# Copper-Mediated Antimicrobial Activity : Targeting Cytochrome *c* Complexes Across Bacterial Species

**DOI:** 10.64898/2026.10.17.759849

**Authors:** Anne Durand, Anne-Soisig Steunou, Monis Athar Khan, Marie-Line Bourbon, Corinne Cassier-Chauvat, Franck Chauvat, Soufian Ouchane

## Abstract

Copper (Cu) exerts potent antibacterial activity and contributes to mammalian immune defense. Although solvent exposed 4Fe-4S clusters in cytoplasmic dehydrogenases are major intracellular targets of Cu, its chemical reactivity and subcellular distribution suggest that additional cellular processes may be vulnerable to Cu toxicity. Here, we identify *c*-type cytochrome biogenesis as a conserved target of Cu excess across diverse pathogenic and environmental bacteria. Analysis of respiratory membrane complexes from pathogens, including *Vibrio cholerae* and *Pseudomonas aeruginosa*, revealed pronounced and selective sensitivity of *c*-type cytochrome-containing complexes to excess Cu. In *P. aeruginosa*, elevated Cu impaired the *cbb_3_*-type cytochrome *c* oxidase and NapBC-containing cytochrome *c* complexes while largely sparing complexes independent of *c*-type cytochromes, including the *aa_3_*-type cytochrome *c* oxidase and succinate dehydrogenase. In contrast, the *caa_3_*-type cytochrome *c* oxidase of *Bacillus subtilis*, which contains a covalently bound cytochrome *c* subunit, was also severely affected. Cu likewise depleted *c*-type cytochromes in plant-associated and photosynthetic bacteria, indicating that this vulnerability is broadly conserved. Complementary analyses in *B. subtilis* and *Escherichia coli* showed that Cu primarily disrupts *c*-type cytochrome assembly rather than promoting the degradation of preassembled complexes. These findings establish *c*-type cytochrome-containing respiratory complexes as conserved targets of Cu toxicity. Given their central role in electron transport and adaptation to oxygen-limited environments, disruption of cytochrome *c* biogenesis may contribute to Cu antimicrobial activity and expose a potential vulnerability in pathogens that rely on *c*-type cytochromes for microaerobic respiration and colonization of hypoxic host tissues.

## Introduction

*c*-Type cytochromes are widespread bacterial electron carriers that participate in diverse energy producing pathways, including photosynthesis and both aerobic and anaerobic respiration [1]. These proteins carry covalently attached heme groups, linked through thioether bonds formed between the heme vinyl groups and the cysteine residues of the conserved CXXCH heme-binding motif in the apocytochrome, with the histidine residue serving as an axial ligand to the heme iron. In Gram-negative bacteria, *c*-type cytochrome assembly occurs in the periplasm, whereas in Gram-positive bacteria it takes place at the extracytoplasmic side of the membrane [2]. A critical step in this process is the formation of thioether bonds, which requires the cysteine thiols of the heme-binding motif to remain reduced prior to heme attachment. Accordingly, disruption of thiol-reduction pathways severely impairs cytochrome *c* maturation, indicating that oxidative conditions in the periplasm are likely to interfere with cytochrome *c* assembly. *c*-type cytochromes are key components of diverse respiratory pathways, including the *cbb_3_*-type cytochrome *c* oxidase, a terminal oxidase that enables efficient respiration under oxygen-limited conditions [3]. They have been implicated in pathogenesis and tissue invasion across diverse bacterial pathogens, including *Pseudomonas (P.) aeruginosa* [4], *Escherichia (E.) coli* [5,6]*, Vibrio (V.) cholerae* [7], *Mycobacterium (M.) tuberculosis* [8]*, Salmonella (S.) Typhimurium* [9]*, Neisseria (N.) gonorrhoeae* [10]*, Staphylococcus (S.) aureus* [11] and *Klebsiella (K.) pneumoniae* [12]. In the opportunistic pathogen *P. aeruginosa*, the electron transport chain is similarly linked to host tissue invasion and adaptation to stressful conditions, with the *cbb_3_*-type cytochrome oxidase contributing to biofilm formation and virulence [4]. In *V. cholerae*, maintaining functional terminal oxidases *bd* and *cbb_3_* has likewise been shown to be essential for establishing infection and proliferating in the mouse intestine [7]. Given this central role in respiration and virulence, factors that compromise *c*-type cytochrome biogenesis or function have the potential to impair bacterial fitness and pathogenicity. Many respiratory and metabolic pathways including photosynthesis, depend on transition metals such as iron (Fe) and copper (Cu), which serve as essential redox cofactors [13]. However, their redox activity and high affinity for biological ligands also make metals toxic. This property is exploited by the mammalian immune defense system [14,15]. During infection, the host actively redistributes Cu to pathogen exposed sites, including the gastrointestinal and respiratory tracts, the bloodstream, and immune cell compartments. Phagocytic cells can further accumulate Cu within maturing phagolysosomes, where elevated Cu levels promote microbial killing by disrupting essential cellular processes in engulfed pathogens [14,15]. The ability of bacteria to withstand excess Cu depends on the expression of copper homeostasis systems. In most bacteria, Cu tolerance involves the induction of at least the Cu(I)-efflux ATPase CopA. In a limited number of species, including *E. coli, P. aeruginosa,* and *K. pneumoniae*, Cu efflux is also mediated by the CusABC complex [16–19]. Several studies have demonstrated that these Cu-efflux systems contribute to bacterial virulence and promote survival during infection by enabling pathogens to withstand Cu-mediated host innate immune defenses [17,20–22].

Cu, particularly in its reduced Cu(I) form, is extremely reactive even at low concentrations. Decades of research have provided evidence that Cu excess causes cellular damage across all domains of life, through mechanisms including reactive oxygen species production, DNA damage, protein mis-metallation, and disruption of iron-sulfur cluster proteins and their biogenesis [23]. In bacteria, however, most characterised mechanisms of Cu toxicity involve cytoplasmic targets in striking contrast to the localisation of bacterial cuproproteins, which are predominantly found in the inner membrane and periplasm, implying active Cu trafficking across these compartments. Because Cu(I) is highly redox-active and displays strong affinity for sulfur-containing residues, it can promote thiol oxidation and catalyse disulfide bond formation in the bacterial periplasm [24,25], suggesting that sulfur-containing proteins in this compartment may represent particularly sensitive targets of Cu toxicity. Consistently, using a CopI-deficient mutant that accumulates Cu(I) within the periplasm, we showed that Cu(I) selectively impairs membrane-associated cytochrome *c* oxidase activity by affecting *c*-type cytochromes, identifying cytochrome *c* biogenesis as a potential target of Cu toxicity [26]. In contrast, mutants lacking the CopA Cu-efflux ATPase accumulate Cu(II) primarily in the cytoplasm, where respiratory defects arise indirectly through inhibition of heme biosynthesis caused by Cu-mediated damage to the radical SAM enzyme coproporphyrinogen III dehydrogenase (HemN) [27]. Similar observations have been reported in *N. gonorrhoeae* and are further supported by *in vitro* studies demonstrating the susceptibility of exposed 4Fe-4S clusters in radical SAM enzymes to Cu-mediated damage [10,27,28]. Together, these observations suggest that the cellular consequences of Cu toxicity depend strongly on the compartment in which Cu accumulates.

Because (i) known bacterial cuproproteins are localised mainly in the periplasm and inner membrane, (ii) excess environmental copper first accumulates within the periplasm, and (iii) Cu-efflux systems export excess cytoplasmic Cu(I) into the periplasm, Cu overload is expected to occur primarily in extracytoplasmic compartments. We therefore hypothesised that the bacterial membrane and periplasm constitute major sites of Cu toxicity, and that bacteria relying on *c*-type cytochromes would be particularly sensitive to Cu excess. To test this hypothesis, we examined a broad range of environmental and pathogenic bacteria with distinct respiratory lifestyles. Our findings support a general mechanism whereby Cu(I) selectively disrupts *c*-type cytochromes within the periplasm and membrane, confirming cytochrome *c* biogenesis as a broadly conserved target of copper toxicity.

## Results

### Cu excess decelerates respiration by selectively impairing the cytochrome *c* oxidase-dependent respiratory branch

The respiratory electron transport chain in *Rubrivivax (R.) gelatinosus* branches downstream of the quinone pool [29] into two alternative pathways: a cytochrome *c*-independent route involving the *bd*-type ubiquinol oxidase, and a cytochrome *c*-dependent pathway in which electrons are transferred via the *bc*_1_ complex to the *cbb*_3_ cytochrome *c* oxidase (**Fig. 1A**). Given that excess Cu(I) has been reported to impair *c*-type cytochromes in the *R. gelatinosus* ΔcopI mutant, which accumulates Cu(I) in the periplasm of *R. gelatinosus* [26], we hypothesized that the aerobic cytochrome *c*-dependent respiratory chain may be compromised under Cu overload.

**Figure 1A:**
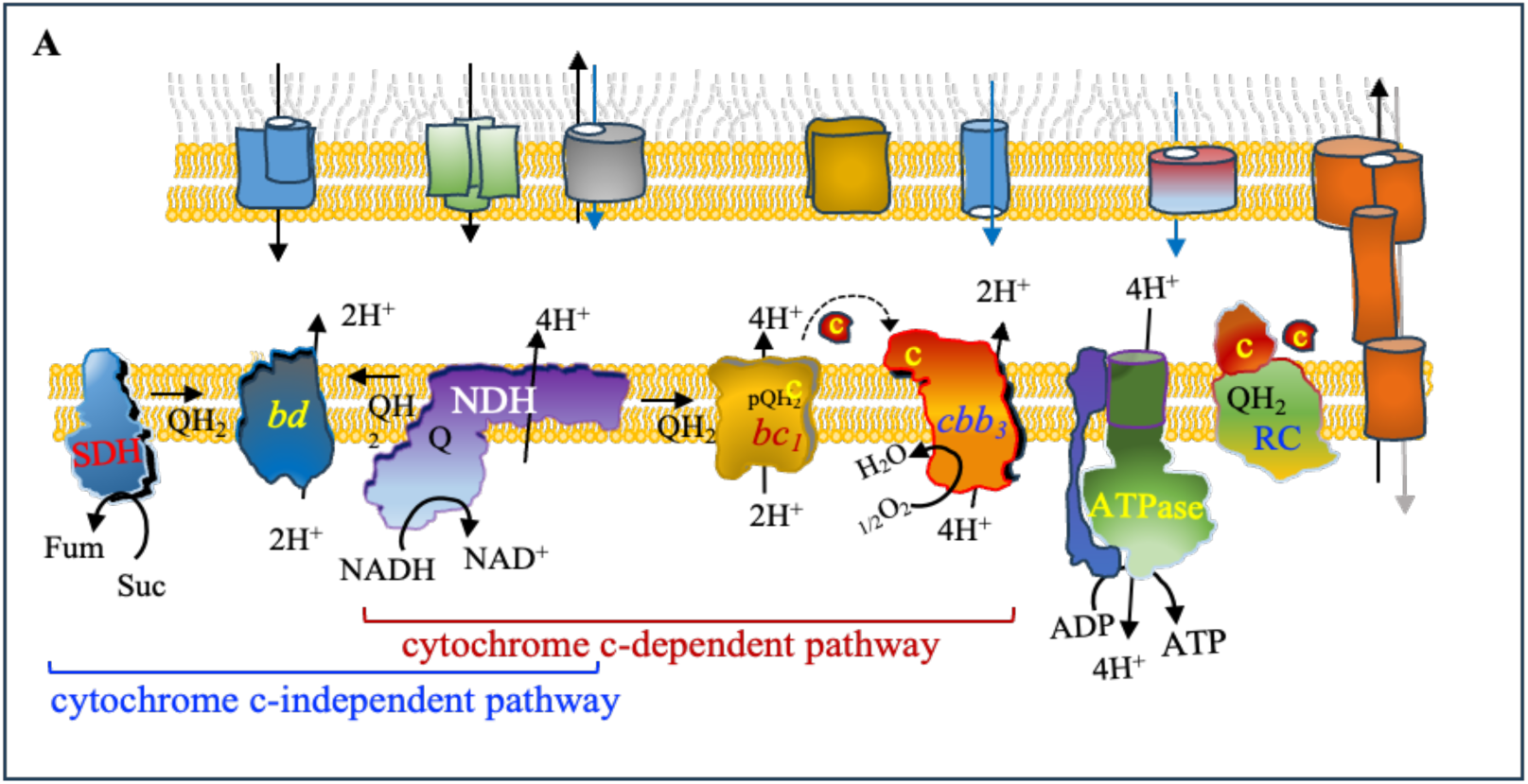
Respiratory electron transport pathways and the effect of CuSO_4_ on aerobic respiration in *R. gelatinosus*. **A-**Schematic representation of the *R. gelatinosus* cytoplasmic membrane, illustrating the branched respiratory electron transport chain. The cytochrome *c*-independent pathway involves the *bd*-type ubiquinol oxidase, whereas the cytochrome *c*-dependent pathway involves electron transfer through the cytochrome *bc_1_* complex to the *cbb*_3_-type cytochrome *c* oxidase. Complexes depicted in the outer membrane represent putative transporters. SDH, succinate dehydrogenase; NDH, NADH dehydrogenase; *bc_1_*, cytochrome *bc_1_* complex; *cbb*_3_, cytochrome *c* oxidase; RC, photosynthetic reaction center.

To evaluate the impact of Cu(I) on each respiratory branch, we analyzed 120 strains selectively relying on either the *bc*1 / *cbb*3 type cytochrome oxidase pathway or the *bd* quinol oxidase pathway in the presence or absence of 1000 μM added CuSO4. Dissolved oxygen consumption were monitored in wild-type (WT), *cbb*3 deficient, and *bd-*deficient strains [29] grown semi aerobically in malate medium,, with or without 1000 μM CuSO4, using a real-time respiration assay (PreSens). In WT cells, growth in the presence of 1000 μM CuSO4 caused only a moderate decrease in oxygen consumption rate (**Fig. 1B**). In contrast, the *cbb*3 (*cbb*3-) deficient mutant, which relies exclusively on the *bd*-type quinol oxidase branch, showed no detectable alteration in oxygen consumption upon exposure to of CuSO4, indicating that this respiratory pathway and the *bd* quinol oxidase are not affected by Cu(I). Conversely, CuSO4 markedly decreased the rate of respiration in the *bd* (*bd*-) deficient mutant, which depends on the *bc*1 complex and *cbb*3 cytochrome *c* oxidase for oxygen reduction (**Fig. 1B**).

**Figure 1B:**
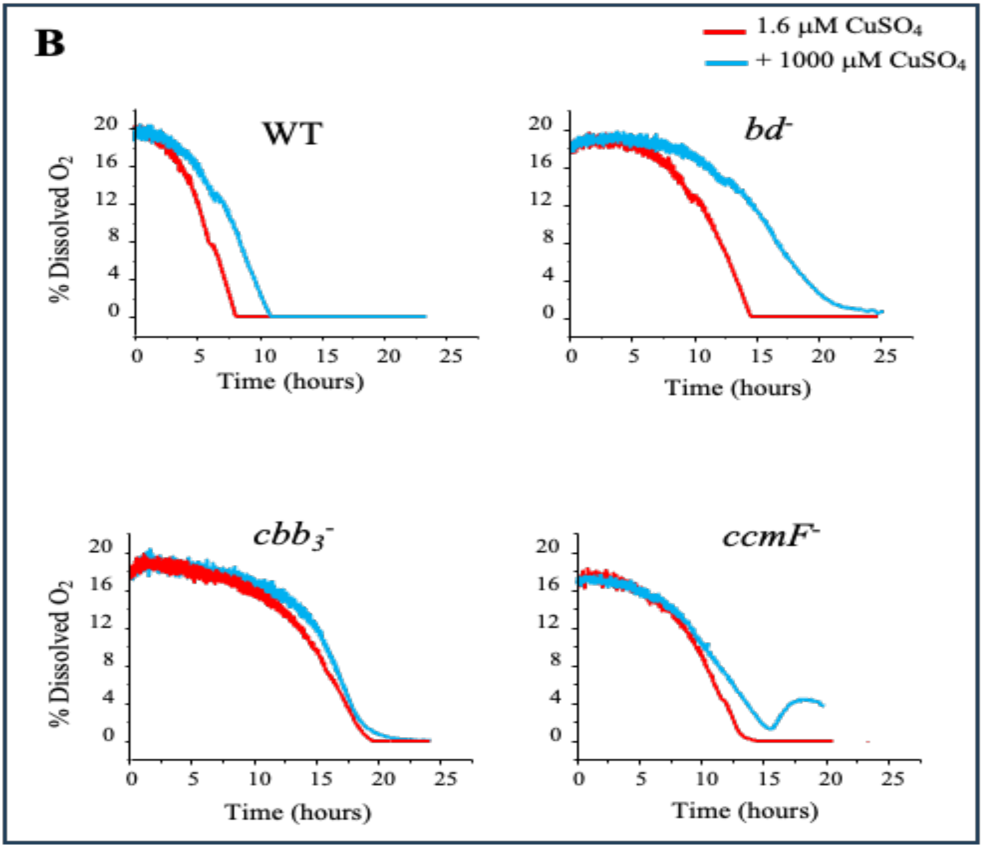
Effect of CuSO_4_ on aerobic respiration: Real-time oxygen consumption was measured in WT, *cbb*_3_^-^, *bd*^-^ and *ccmF*^-^mutant cells grown semi-aerobically in medium supplemented with or without 1000 μM CuSO_4_.

In *R. gelatinosus*, the biogenesis of *c*-type cytochromes depends on the Ccm machinery, which mediates heme attachment to periplasmically translocated apocytochromes *c*. A ΔccmF mutant previously shown to lack detectable levels of cytochrome *c* [26] was therefore used as an additional control to evaluate the impact of Cu on respiration. Similar to the *cbb_3_* mutant, the ΔccmF strain which relies exclusively on the quinol oxidase branch and showed no significant change in oxygen consumption in the presence of CuSO_4_ (**Fig. 1B**). Together, these results demonstrate that (i) the *bd* quinol oxidase can sustain oxygen reduction and supports aerobic growth under conditions of Cu stress, whereas excess Cu specifically affects the cytochrome *c-*dependent branch of the respiratory electron transport chain. Furthermore, the pronounced Cu sensitivity of the *bd-*deficient strain suggest that *cbb_3_* cytochrome *c* oxidase is a major target of Cu toxicity in the respiratory chain of Cu-overloaded *R. gelatinosus* cells.

### Cu-induced decrease in respiration correlates with a decrease in cytochrome *c* oxidase complexes but not in succinate dehydrogenase

As an initial step toward understanding how excess Cu affects respiration, we previously employed a ΔcopI mutant to increase periplasmic Cu(I) levels, and demonstrated that cytochrome *c* assembly is impaired in this genetic background [26]. In *R. gelatinosus*, the Cu-efflux pump CopA exports cytoplasmic Cu(I) into the periplasm. Since this organism lacks a Cus system, we anticipated that excess Cu may result in transient Cu(I) accumulation in the periplasm, thereby affecting periplasmic processes. To identify bioenergetic complexes affected by elevated periplasmic Cu(I), WT cells were grown in the presence of increasing concentrations of CuSO_4_ (1.6 to 1600 μM) and equal amounts of membrane proteins were solubilized and analyzed by blue native PAGE (BN-PAGE). As shown in **Fig. 2A**, the abundance of the tetraheme cytochrome *c*-containing reaction center (RC) decreased markedly upon exposure to 400 μM CuSO_4_. In contrast, *in-gel* activity assays for quinone oxidoreductases: NADH dehydrogenase (NDH) and succinate dehydrogenase (SDH) showed that both complexes remained fully active even at high (1600 μM) CuSO_4_ concentration, indicating that these complexes are resistant to Cu stress. In contrast, the *cbb*_3_ cytochrome *c* oxidase activity in the same samples showed a pronounced decrease at 200 μM CuSO_4_ and an almost complete loss of activity at 400 μM CuSO_4_ (**Fig. 2A**).

**Figure 2A:**
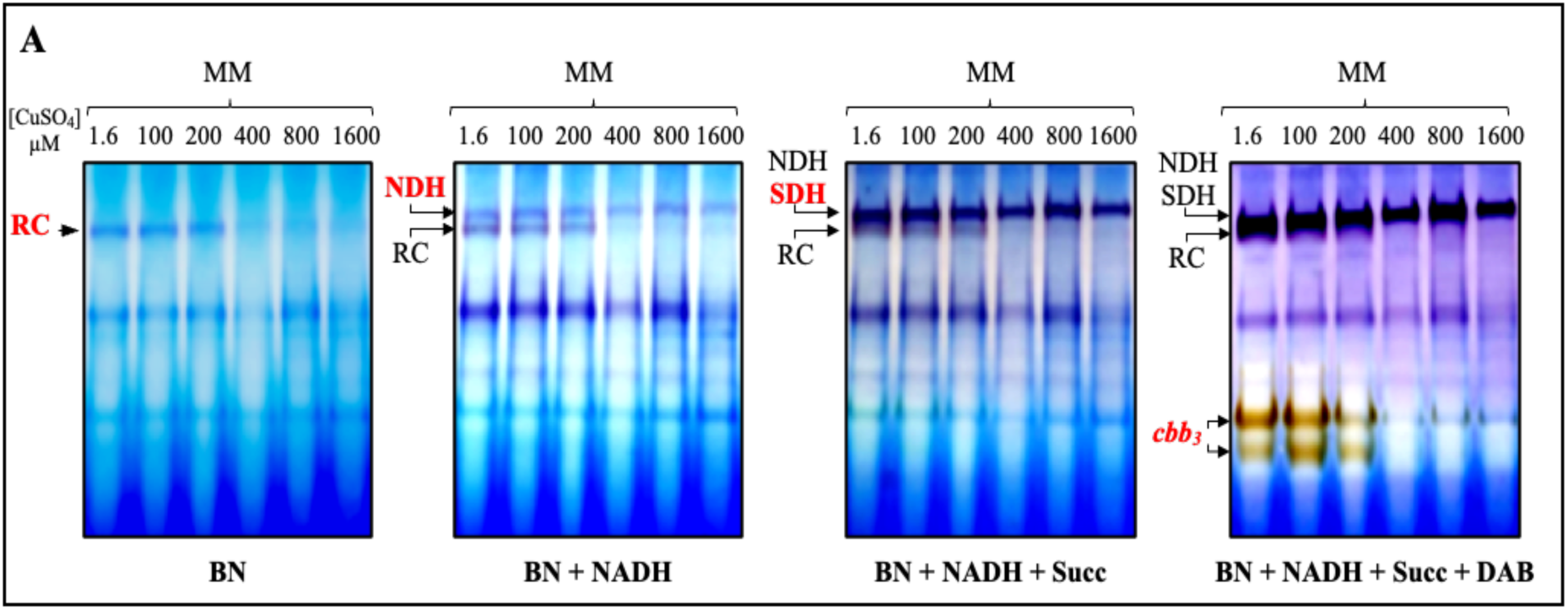
Cu overload selectively disrupts *c*-type cytochromes in the *R. gelatinosus* respiratory chain: **A-**Cells were grown in MM supplemented with increasing concentration of CuSO_4_ (0 - 1600 μM). Solubilized membranes were separated by BN-PAGE and sequentially assayed for NADH dehydrogenase, succinate dehydrogenase (SDH), and *cbb*_3_ oxidase activities.

Collectively, these results indicate that respiratory complexes containing *c*-type cytochromes are particularly sensitive to Cu(I), whereas SDH, which contains only cytochrome *b*-type, and NDH which contains flavin mononucleotide cofactor and unexposed iron-sulfur clusters, are insensitive to increasing Cu(I) stress. These observations were further corroborated by reduced minus oxidized difference spectra of membranes exposed to increasing CuSO_4_ concentrations. The spectra revealed a progressive decrease in *c*-type cytochromes as indicated by the peak at 552 nm for cells exposed to increasing Cu concentration, whereas the cytochrome *b* content (peak at 560 nm) remained largely unchanged (**Fig. 2B**). Consistent with these results, tetramethylbenzidine (TMBZ) based heme staining confirmed a global reduction of membrane-associated *c*-type heme proteins, including the tetraheme cytochrome *c* subunit of the reaction center (PufC), cytochrome *c* peroxidase (CcpA), both CcoO and CcoP *c*-type cytochrome subunits of *cbb*_3_ cytochrome *c* oxidase, and the cytochrome *c*′ (**Fig. 2C**).

**Figure 2B-C:**
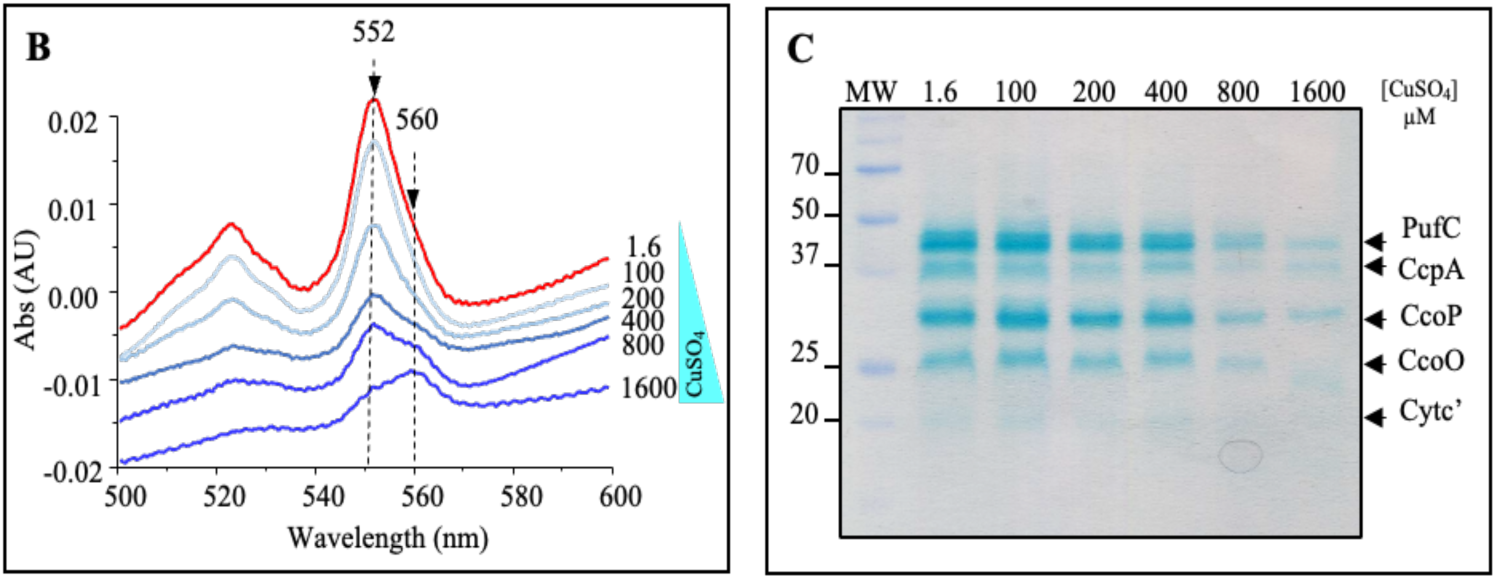
Cu overload selectively disrupts *c*-type cytochromes in the *R. gelatinosus* respiratory chain: **B-** The specific decrease in the characteristic peak at 552 nm indicated a reduction in *c*-type cytochromes content. **C-**Detection of *c*-type cytochromes by heme-staining. Equal amounts of membrane proteins were separated on 12% SDS-PAGE and *c* cytochromes were visualized by heme-staining in the presence of TMBZ. CcoO and CcoP are subunits of *cbb_3_* cytochrome *c* oxidase, PufC to the tetraheme cytochrome of RC. CcpA corresponds to the cytochrome *c* peroxidase and c’, corresponds to the soluble cytochrome *c*.

### CopI is required to maintain periplasmic cytochrome c biogenesis under copper stress in Vibrio cholerae

In the human intestinal pathogen *Vibrio (V.) cholerae* O1 El Tor, CopI (VCA0261–0260) has previously been implicated in periplasmic Cu(I) resistance, similarly to its role in *R. gelatinosus* [30,31]. CopI may therefore contribute to the protection of cytochrome *c* assembly and *cbb*_3_ cytochrome *c* oxidase activity. To test this hypothesis, a *copI* mutant (*copI*::*Tn*) of *V. cholerae* [32] was grown in LB medium or LB supplemented with 150, 250, or 500 μM CuSO_4_, and the relative abundance of cytochrome *c* and *b*-type hemes was assessed by reduced minus oxidized difference spectroscopy of whole cell extracts. As shown in **Fig. 3A**, Cu exposure resulted in a progressive decrease in *c*-type cytochromes, whereas *b*-type cytochromes remained largely unaffected. Compared with cells grown in LB alone, *copI*::*Tn* cells exposed to CuSO_4_ showed a marked reduction in *c*-type cytochromes at 150 μM and a complete loss at 250 and 500 μM CuSO_4_, whereas cytochrome *b* signals persisted. In contrast, reduced-minus-oxidized spectra showed that WT cells retained detectable levels of *c*-type cytochromes even at 250 and 500 μM CuSO_4_ (**Fig. 3B**). Yet the amount of *c*-type cytochromes decreased at 500 μM CuSO_4_, and was substantially reduced at 1000 μM CuSO_4_ in the WT. We next examined whether this protection extended to *cbb_3_* cytochrome *c* oxidase activity. *In-gel* activity assays following BN-PAGE analysis of membrane complexes showed a strong decrease in *cbb_3_* cytochrome *c* oxidase activity at 500 μM CuSO_4_, with only trace activity remaining, while the activity was completely abolished at 1000 μM CuSO_4_ (**Fig. 3C**). At 250 μM CuSO_4_, the activity of *cbb_3_* was increased likely due to the higher availability of Cu required for oxidase assembly. Together, these findings establish that CopI is essential for maintaining periplasmic cytochrome *c* biogenesis in *V. cholerae* under Cu stress and indicate that excess Cu(I) compromises *c*-type cytochrome biogenesis in this organism.

**Figure 3:**
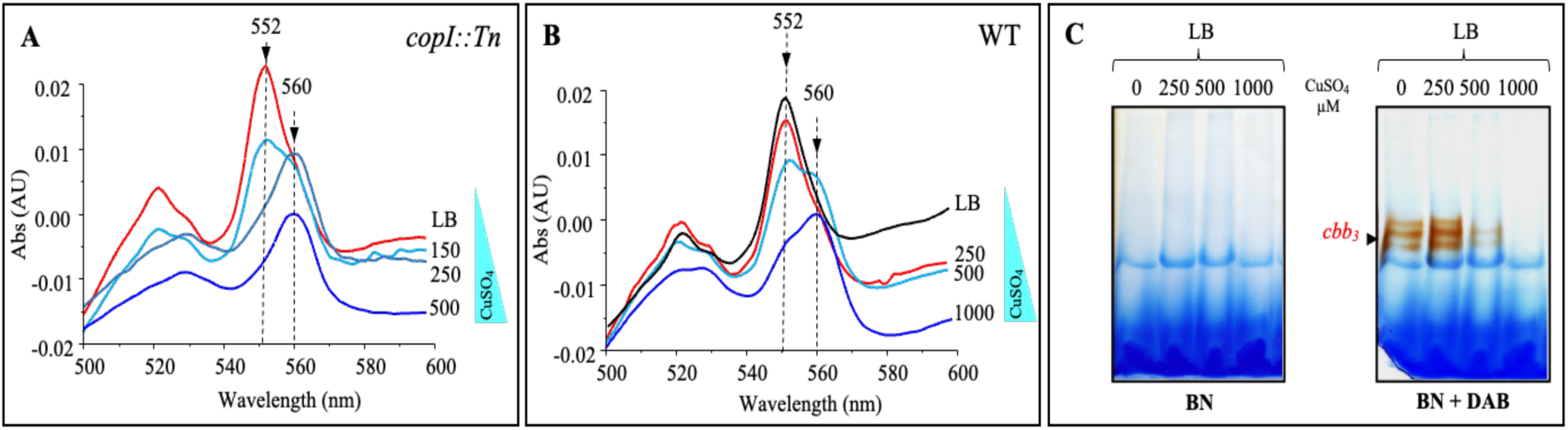
Cu excess depletes *c*-type cytochromes and impairs *cbb_3_* cytochrome *c* oxidase in *Vibrio cholerae*. **A-**Reduced minus oxidized difference absorption spectra of membranes isolated from *copI::Tn* mutant cells grown in medium supplemented with increasing CuSO4 concentrations (150 - 500 μM). The substantial decrease in the characteristic absorption peak at 552 nm indicates a reduction in *c*-type cytochrome content. **B-**Reduced-minus-oxidized difference absorption spectra of membranes isolated from WT cells grown in medium supplemented with increasing concentrations of CuSO_4_ (250 to 1000 μM). **C-**Solubilized membranes were separated by BN-PAGE and assayed for *cbb*_3_ oxidase activity.

### Respiratory chain plasticity contributes to periplasmic Cu resistance in P. aeruginosa

*Pseudomonas aeruginosa*, a major opportunistic human pathogen, possesses several Cu-resistance mechanisms contributing to the its survival in Cu-rich environments [33]. Recently, Cu(I) was shown to induce the DsbC, DsbD/DsbE reductive pathway, which repair aberrantly formed disulfide bonds in the periplasm of *P. aeruginosa* [33,34]. Because oxidized thiols of cytochrome *c* precursors are substrates of this reductive pathway, we investigated the impact of Cu excess on cytochrome *c* biogenesis in *P. aeruginosa* PAO1 by monitoring changes in the cellular cytochrome *c*-type to *b*-type ratio. WT cells were grown in LB medium supplemented with increasing concentrations of CuSO_4_ (200 - 1600 μM), and reduced minus oxidized difference spectra were recorded from whole cell extracts. As observed in *R. gelatinosus*, increasing Cu concentrations caused a progressive and marked decrease in the 552 nm absorption peak characteristic of *c*-type cytochromes in *P. aeruginosa*, whereas the 560 nm peak corresponding to *b*-type cytochromes remained largely unchanged (**Fig. 4A**). These data indicate a substantial reduction in *c*-type cytochrome content in Cu-exposed cells. To determine how excess Cu(I) affects the respiratory chain, membrane proteins were solubilized and analyzed on BN-PAGE. *In-gel* cytochrome *c* oxidase activity staining revealed that the *cbb_3_* cytochrome *c* oxidase remained active up to 800 μM CuSO_4_ and even showed increased activity with rising copper concentrations before declining at 1000 μM CuSO_4_ (**Fig. 4B**). Notably, the decrease in *cbb_3_* activity was accompanied by a pronounced increase in *aa_3_* cytochrome *c* oxidase activity. Unlike the *cbb_3_* complex, which contains two membrane-bound *c*-type cytochrome subunits essential for its assembly and function, the *aa_3_* oxidase lacks *c*-type cytochromes, likely rendering it non-susceptible to Cu(I) mediated damage. BN-PAGE gels were subsequently incubated with succinate to assess succinate dehydrogenase (SDH) activity. Although SDH activity was lower in the control, it remained fully preserved across all increased CuSO_4_ concentrations tested. An additional succinate-dependent complex (Cytc^+^) was detected on BN-PAGE and displayed marked sensitivity to Cu, with its activity substantially decreasing at 800 μM CuSO_4_ and disappearing completely at 1600 μM CuSO_4_ (**Fig. 4B**).

**Figure 4A:**
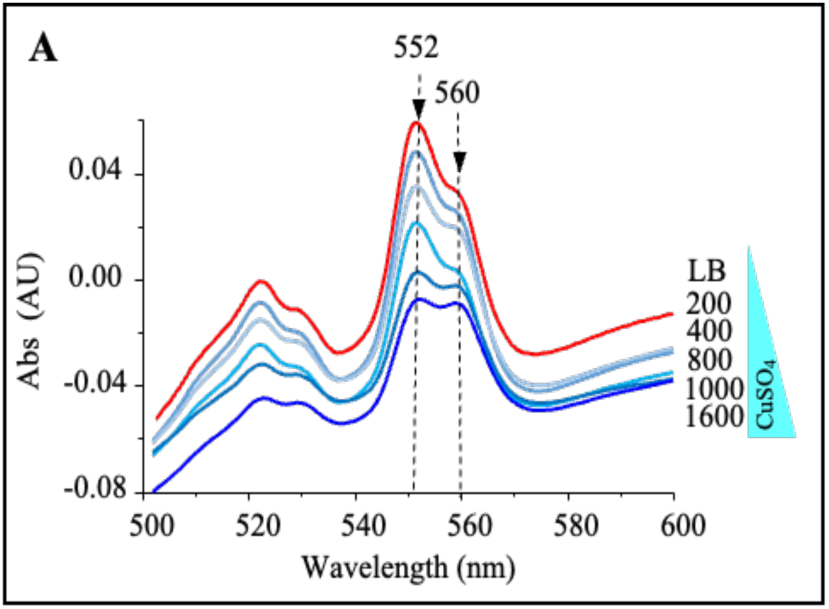
*cbb_3_* Oxidase activity and *c*-type cytochrome biogenesis are sensitive to Cu stress in *Pseudomonas aeruginosa.* **A-**Reduced-minus-oxidized difference absorption spectra of membranes isolated from WT cells grown in medium supplemented with increasing concentrations of CuSO_4_ (200 to 1600 μM). The decrease in the peak at 552 nm indicates a reduction in *c*-type cytochrome content.

The progressive loss of the signal intensity strongly suggested the presence of one or more *c*-type cytochromes within this complex. To verify the presence of cytochrome *c* subunit(s) in this complex, the Cytc⁺ active band was excised from the stained BN-PAGE, resolved in a second dimension by SDS-PAGE followed by TMBZ staining for cyt*c* detection. The SDH and *aa_3_* bands were analyzed in parallel as negative staining controls. Two heme-containing subunits of approximately 15 and 25 kDa were detected in the Cytc⁺ band, whereas no corresponding bands were observed in the controls (Fig. S1). A genomic analysis of *P. aeruginosa* focusing on *c*-type cytochromes suggested that the detected bands 15 and 25 kDa bands could correspond to NapB and NapC, the two cytochromes *c* containing subunits of the periplasmic nitrate reductase complex (NapABCDE) in *P. aeruginosa* [35]. Notably, a succinate:quinone oxidoreductase:nitrate reductase supercomplex was identified in *B. subtilis* [36] based on the colocalization of succinate:NBT and MV:nitrate oxidoreductase activities on BN-PAGE, and further supported by LC-MS/MS analysis. The fortuitous detection in *P. aeruginosa* of this *c*-type cytochrome-containing complex in our analysis provides additional support for the Cu (I) sensitivity of *c*-type cytochromes in bacteria.

Finally, TMBZ heme staining confirmed a decrease in all membrane-associated *c*-type heme proteins (**Fig. 4C**), including the two Nap cytochrome *c* subunits, the cytochrome *c* peroxidase CcpA, and the *cbb_3_* cytochrome *c* oxidase subunits CcoO and CcoP. Together, these results indicate that Cu exposure preferentially affects *c*-type cytochromes, while sparing *b* and *a*-type cytochromes and SDH activity, and is accompanied by a shift in terminal oxidase activity from *cbb_3_* toward *aa_3_.* This respiratory chain plasticity and remodeling may significantly contribute to the ability of *P. aeruginosa* to maintain respiratory activity under Cu stress.

**Figure 4B:**
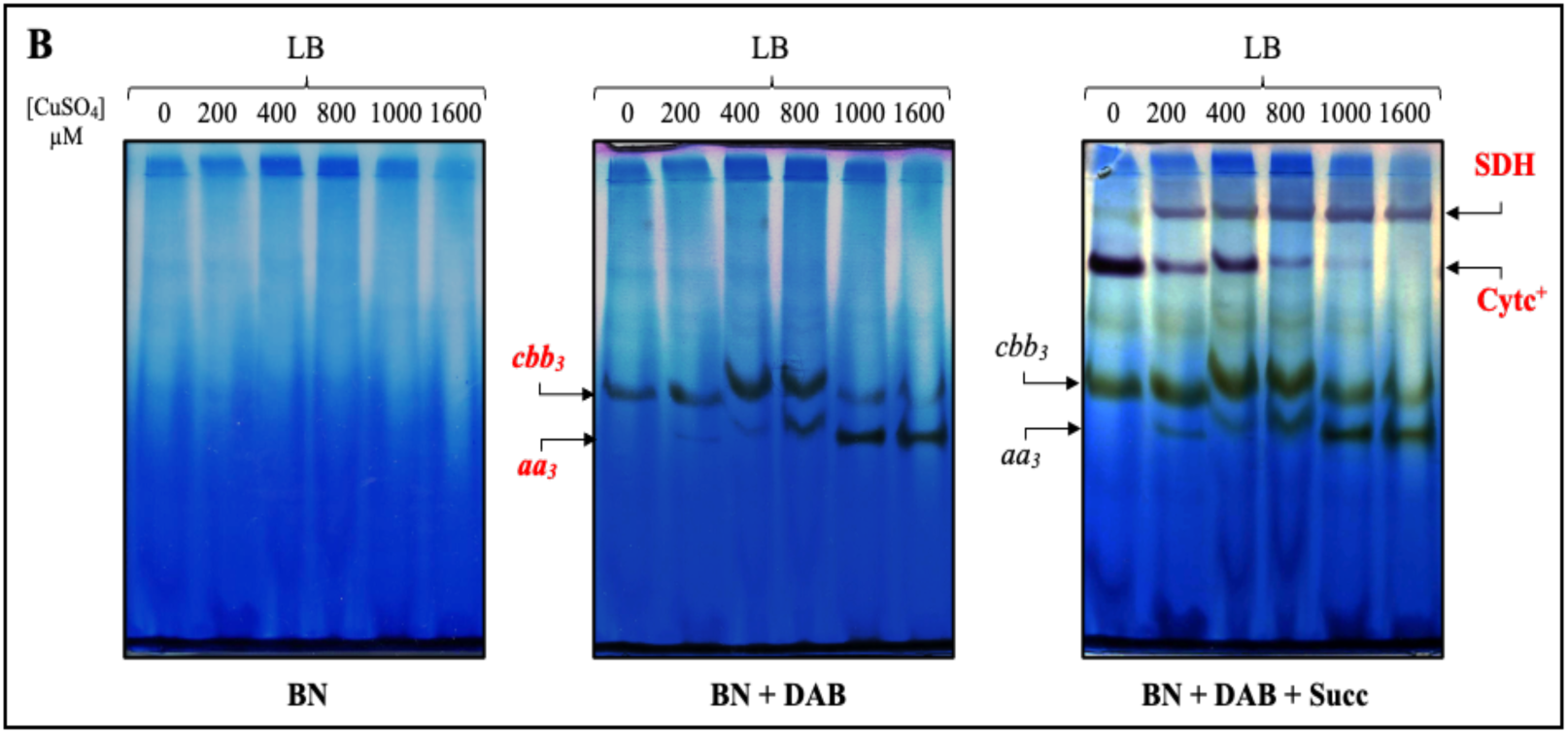
*cbb_3_* Oxidase activity and *c*-type cytochrome biogenesis are sensitive to Cu stress in *Pseudomonas aeruginosa.* Solubilized membrane proteins were separated by BN-PAGE and sequentially assayed for *cbb*_3_ oxidase and succinate dehydrogenase (SDH) activities.

**Figure 4C:**
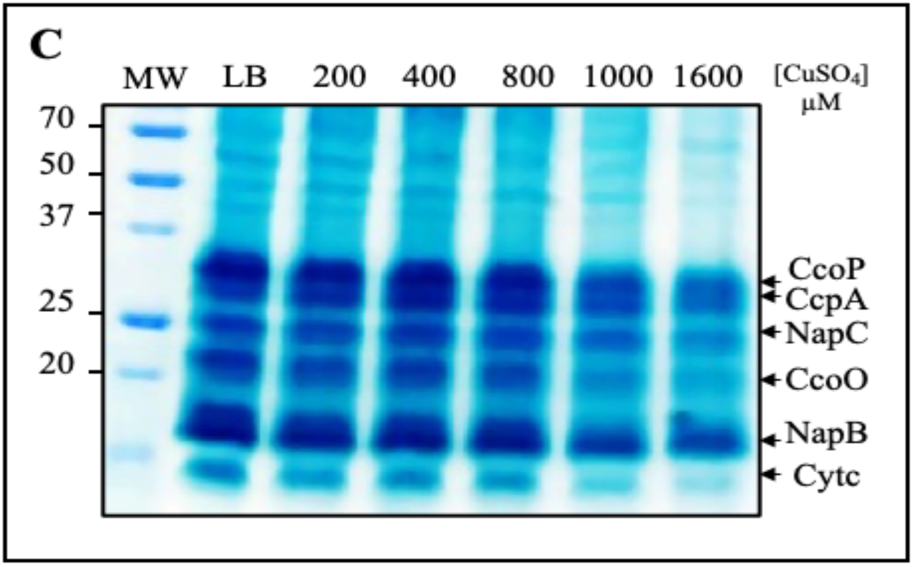
*cbb_3_* Oxidase activity and *c*-type cytochrome biogenesis are sensitive to Cu stress in *Pseudomonas aeruginosa.* Detection of membrane-associated *c*-type cytochromes of *P. aeruginosa.* Equal amounts of membrane proteins were separated on 12% SDS-PAGE, and *c* cytochromes were visualized using TMBZ. CcoO and CcoP are subunits of *cbb_3_*. CcpA corresponds to the cytochrome *c* peroxidase, NapB and NapC are cytochrome *c* subunits of the nitrate reductase. Cyt*c*, corresponds to the soluble cytochrome.

### Evidences from plant-associated bacteria confirm that Cu specifically decreases the of *c*-type cytochromes

Plant-associated bacteria colonize plant tissues where they may establish either beneficial, or pathogenic interactions. Their energy metabolism relies on terminal respiratory oxidases that support ATP production and adaptation to fluctuating oxygen conditions. Copper compounds, including CuSO_4_, are widely used in agriculture as antimicrobial agents to protect plants against bacterial and fungal pathogens [37]. We therefore examined the effects of excess copper on two plant-associated bacteria with contrasting effects on plants, the pathogen *Pseudomonas (P.) syringae* and the symbiotic nitrogen fixer *Bradyrhizobium (B.) japonicum*.

The genome of *P. syringae* pv. tomato DC3000 encodes a putative *cbb_3_* cytochrome *c* oxidase complex (CcoNOQP), which is predicted to contribute to its aerobic respiratory chain alongside a cytochrome *bd* type quinol oxidase and a *bo_3_* type quinol oxidase. Cells were grown in LB medium supplemented with increasing concentrations of CuSO_4_ (250 - 1500 μM), and reduced minus oxidized difference spectra were recorded from whole cell extracts. Unexpectedly, no cytochrome *c*-type signal was detected in the membranes of *P. syringae*; instead, only a peak at 560 nm, characteristic of *b*-type cytochromes, was observed (**Fig. 5A**). Importantly, and in line with our findings, Cu showed no detectable impact on *b*-type cytochromes at any of the concentrations tested, further supporting the notion that Cu(I) selectively affects *c*-type cytochromes. *In-gel* cytochrome *c* oxidase and succinate dehydrogenase activity staining revealed bands, that were not or slightly affected by Cu CuSO_4_ excess (**Fig. 5B**). The absence of cytochrome *c*-type in this strain was further confirmed by TMBZ based heme staining (Fig. S2). A closer examination of the genes involved in cytochrome *c* biogenesis (*ccm*) revealed a frameshift mutation in the *ccmH* gene, resulting in an inactive protein and non-functional CCM machinery, thereby providing a likely explaination for the absence of active *cbb_3_* in *P. syringae*.

**Figure 5:**
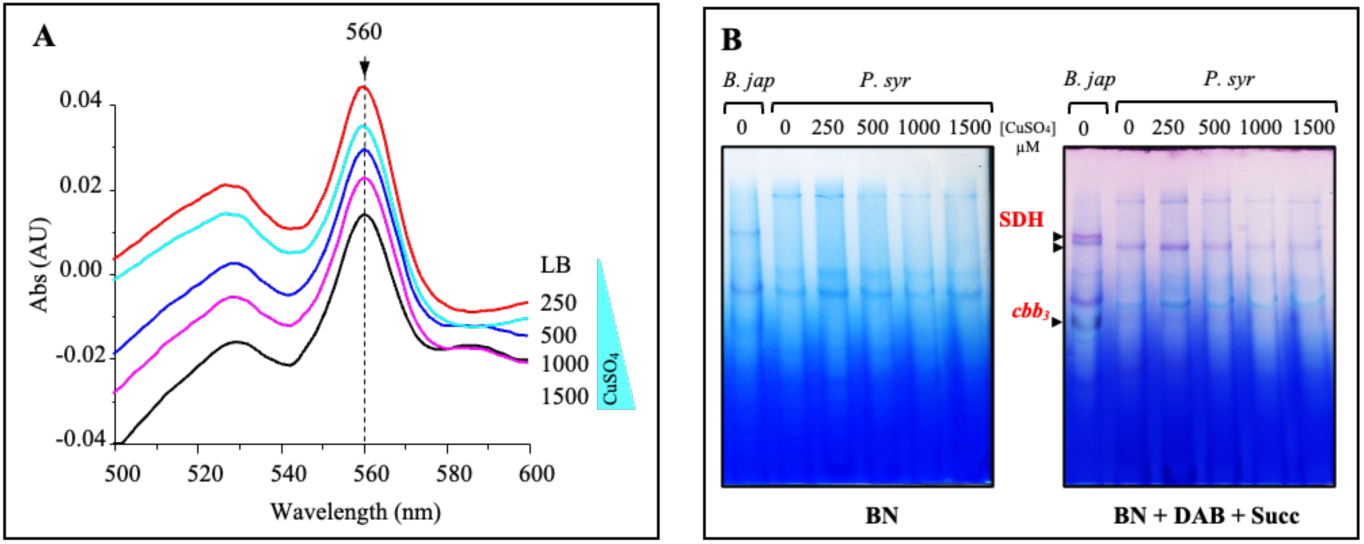
*Pseudomonas syringae* cytochrome *b* complexes withstand Cu stress that disrupts *c*-type cytochromes. **A-**Reduced minus oxidized difference absorption spectra of membranes isolated from *P. syringae* cells grown in medium supplemented with increasing concentrations of CuSO_4_ (250 to 1500 μM) showing no significant change in the amount of cytochrome *b*, as indicated by the characteristic peak at 560 nm. **B-**Solubilized membrane proteins were separated by BN-PAGE and sequentially assayed for *cbb*_3_ oxidase and succinate dehydrogenase (SDH) activities. Membranes isolated from *B. japonicum* were used as positive control for a *cbb_3_* activity.

In contrast *B. japonicum* is known to rely on an active *cbb_3_* cytochrome *c* oxidase for microaerobic respiration and nitrogen fixation [38]. *B. japonicum* was grown in YM medium or YM supplemented with 200, 400, 600 or 1500 μM CuSO_4_, and the relative abundance of cytochrome *c* and *b*-type hemes was analyzed by reduced minus oxidized difference spectroscopy of whole-cell extracts (**Fig. 6A**). Copper exposure led to a decrease in *c*-type cytochromes, whereas *b*-type cytochromes remained unaffected. Furthermore, equal amounts of membrane proteins were solubilized, and the activities of both SDH and *cbb_3_* cytochrome *c* oxidase were analyzed on BN-PAGE (**Fig. 6B**).

**Figure 6:**
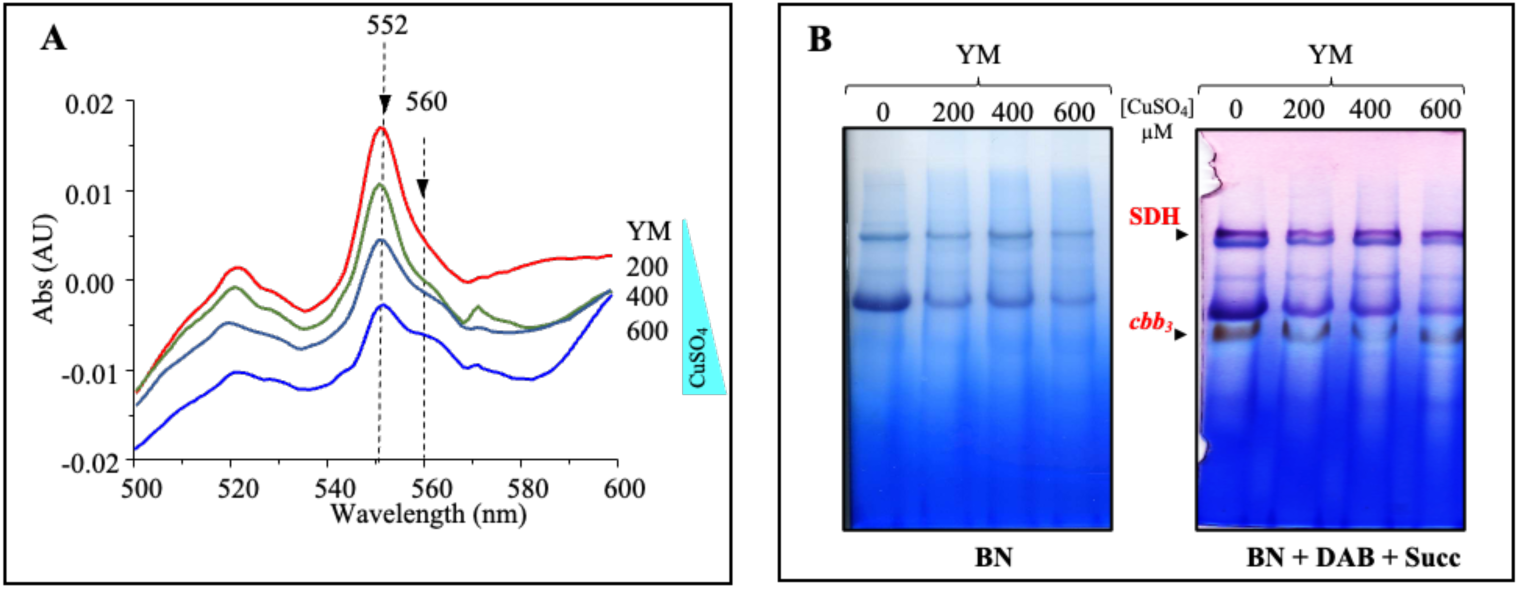
*c*-Type cytochromes and *cbb_3_* oxidase are Cu-sensitive targets in *Bradyrhizobium japonicum*. **A-**Reduced-minus-oxidized difference absorption spectra of membranes isolated from WT cells grown in YM medium supplemented with increasing concentrations of CuSO_4_ (200 to 600 μM). The decrease in the characteristic peak at 552 nm indicates a reduction in cytochrome *c*-type content. **B-**Solubilized membrane proteins were separated by BN-PAGE and sequentially assayed for *cbb*_3_ oxidase and succinate dehydrogenase (SDH) activities.

Consistent with the observations in the other tested strains, exposure to excess Cu selectively reduced the activity of *cbb_3_* cytochrome *c* oxidase, whereas no effect was detected on the heme *b-*type-containing SDH. Taken together, these findings demonstrate that bacteria relying on cytochrome *c*-type oxidases (*cbb_3_*) are particularly susceptible to Cu-mediated growth inhibition. In contrast, bacteria that predominantly depend on *bo_3_* and *bd* quinol oxidases or *aa_3_* cytochrome oxidase maintain their respiratory chain function and are therefore comparatively would be resistant to the detrimental effects of excess copper. These results further raised also the question of whether *c*-type cytochrome containing *caa*_3_ oxidases present in some bacterial species are similarly susceptible to the observed effects of excess copper.

### Threat Posed by Cu on *caa*_3_ oxidase and cytochrome *c* assembly in *Bacillus subtilis*

*Bacillus (B.) subtilis* is a Gram-positive soil bacterium with a branched respiratory chain comprising multiple terminal oxidases. Under aerobic conditions, the cytochrome *bc_1_* complex transfers electrons to the cytochrome *c* oxidase *caa_3_* [39]. This organism therefore provides a suitable model to investigate the effect of Cu on *c*-type cytochromes in a Gram-positive bacterium that relies on the Ccs cytochrome *c* maturation system [40]. To this end, *B. subtilis* was grown in LB medium or LB supplemented with 500, 1000, 1500 or 2000 μM CuSO_4_, and the relative abundance of cytochrome *c, a* and *b*-type hemes was analyzed by reduced minus oxidized difference spectroscopy of whole cell extracts. Cu exposure resulted in a concentration-dependent decrease in both *a*-type (600 nm) and *c*-type (552 nm) cytochromes, whereas *b*-type (560 nm) cytochromes were comparatively less affected (**Fig. 7A**). Then, equal amounts of membrane proteins were solubilized and analyzed by BN-PAGE. The abundance of the active *caa_3_* cytochrome *c* oxidase was markedly decreased upon exposure to 1000 μM CuSO_4_ and was completely abolished in cells grown in the presence of 1500 μM CuSO_4_, consistent with the changes observed in reduced minus oxidized difference spectra (**Fig. 7B**). Notably, super-complexes comprising the *caa_3_* cytochrome *c* oxidase, cytochrome *c* and the *bc_1_* complex have previously been described in *Bacillus subtilis* [36]. Consistently, similar super-complexes *caa_3_-bc_1_* were also detected in our BN-PAGE analyses, and were impaired in abundance / activity upon exposure to excess CuSO_4_. In contrast, succinate dehydrogenase activity remained fully preserved even at the highest CuSO_4_ concentration tested (2000 μM), further supporting the resistance of *b*-type heme-containing complexes to Cu-mediated inhibition, as reported above (**Fig. 7B**). These results were further corroborated by TMBZ heme staining analysis of total protein extracts (**Fig. 7C**). Cu exposure caused a progressive decrease in the abundance of the cytochrome *c*-containing *caa_3_* oxidase subunit CtaC, together with a drastic reduction in the soluble cytochrome *c* CccAB. In contrast, the cytochromes associated with the *bc_1_* complex, QcrC and QcrB, were comparatively less affected by copper exposure.

**Figure 7.**
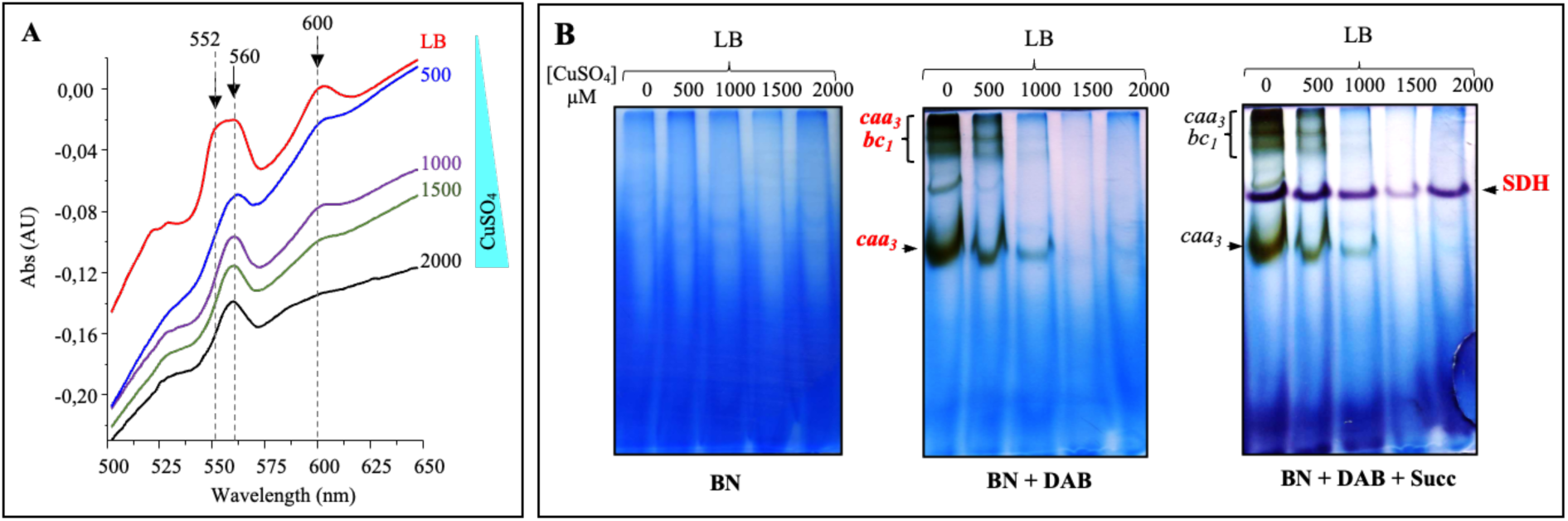
Cu excess depletes *c*-type and *a*-type cytochromes and impairs *caa_3_* cytochrome *c* oxidase activity in *Bacillus subtilis.* **A-**Reduced-minus-oxidized difference absorption spectra of membranes isolated from WT *B. subtilis* cells grown in medium supplemented with increasing concentrations of CuSO_4_ (500 to 2000 μM). The decrease in the characteristic peaks at 552 and 600 nm indicate reductions in cytochrome *c*-type and *a*-type contents, respectively. **B-**Solubilized membrane proteins were separated by BN-PAGE and sequentially assayed for *caa*_3_ oxidase and succinate dehydrogenase (SDH) activities. Putative *caa*_3_::*bc_1_* super-complexes are indicated.

**Figure 7C.**
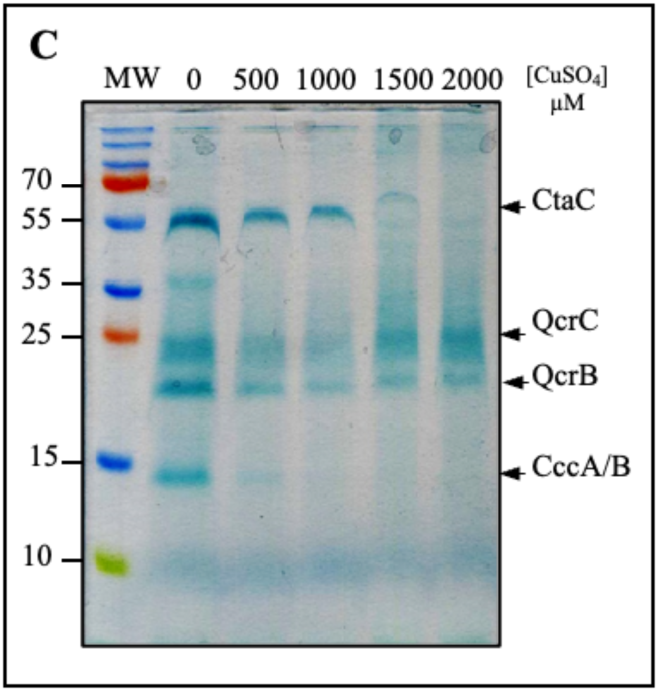
Cu excess depletes *c*-type and *a*-type cytochromes and impairs *caa_3_* cytochrome *c* oxidase activity in *Bacillus subtilis.* Detection of *c*-type cytochrome profiles by heme staining. Equal amounts of membrane proteins were separated on 12% SDS-PAGE, and *c*-type cytochromes were visualized using TMBZ. CtaC corresponds to the cytochrome *c* subunit of *caa_3_*. QcrC and QcrB corresponds to the subunits of cytochrome *bc_1_*, CccA/B corresponds to the soluble cytochromes c_550_ and c_551_ [41].

Taken together, these results demonstrate that excess copper impairs cytochrome *c* complexes, including the *caa_3_* cytochrome oxidase, in *B. subtilis*, and suggest that a similar vulnerability to Cu may extend to other Gram-positive bacteria.

### Copper impairs *c*-type cytochrome assembly rather than degradation of preassembled complexes in *Bacillus subtilis*

Cu can disrupt protein complexes through multiple mechanisms, including protein misfolding, mis-metallation, cofactor displacement, and proteolytic degradation [23]. Owing to its high affinity for thiol containing ligands, copper can interfere with metalloprotein biogenesis and compromise protein assembly or stability. Based on these observations, we previously proposed that excess Cu(I) impairs the assembly of bacterial hemoproteins, particularly cytochrome *c* complexes. However, an alternative possibility is that Cu(I) excess promote the destabilization, and degradation of preassembled cytochrome *c*. To distinguish between these two possibilities, we used *B. subtilis* to compare the effects of continuous CuSO_4_ exposure initiated at the onset of growth with those of an acute CuSO_4_ challenge imposed during stationary growth phase in which overnight grown cells were exposed to CuSO_4_ for 8h. This approach enabled us to differentiate defects in cytochrome *c* biogenesis from Cu-induced destabilization or degradation of preexisting complexes. Reduced minus oxidized difference spectra revealed that continuous exposure to CuSO_4_ caused a progressive decrease in both *a* and *c*-type cytochromes (**Fig. 8A**), consistent with our findings described above. TMBZ heme staining further confirmed decreased levels of the *c*-type heme proteins CtaC and CccAB, whereas the *bc*_1_ complex subunits QcrC and QcrB were only mildly affected or remained unchanged upon Cu exposure, respectively. In striking contrast, acute CuSO_4_ shock of stationary phase cells produced no detectable changes in cytochrome content, irrespective of the CuSO_4_ concentration tested, as assessed by both reduced minus oxidized spectroscopy. Collectively, these findings argue against Cu-induced disassembly or degradation of preexisting *c*-type cytochromes and instead support a model in which excess Cu interferes primarily with cytochrome *c* assembly.

**Figure 8.**
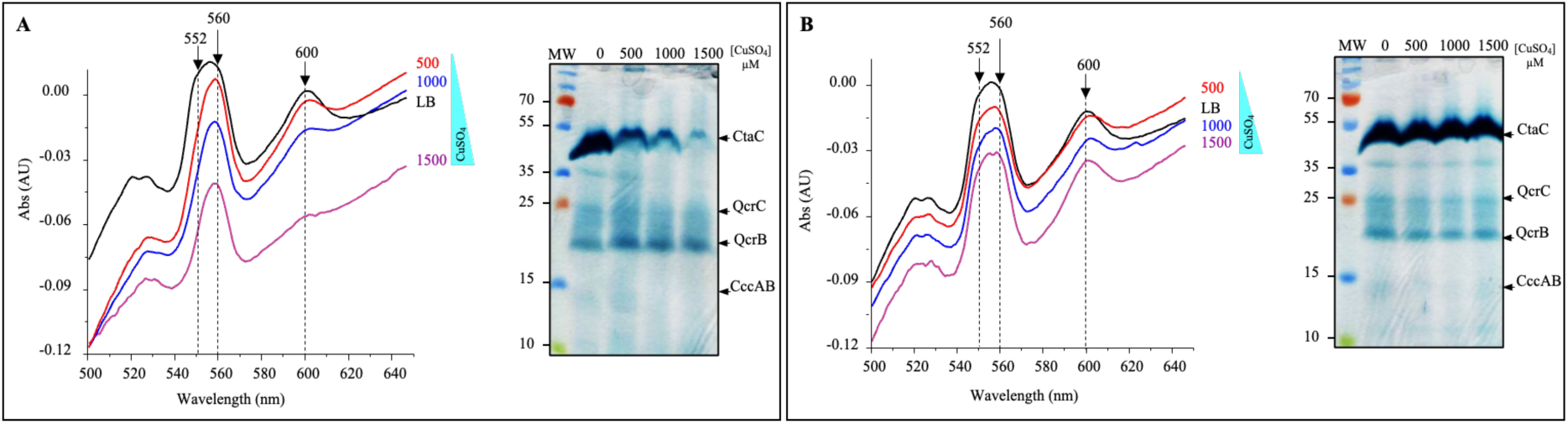
Sustained, but not acute Cu exposure depletes *c*-type and *a*-type cytochromes in *B. subtilis*, reflecting impaired biogenesis rather than degradation. Cells were either grown continuously in LB medium supplemented with increasing concentrations of CuSO_4_ (500 - 1500 μM) or grown overnight in normal LB medium and subsequently exposed to CuSO_4_ for 8 h. (**A**) Reduced minus oxidized difference absorption spectra and heme staining of *c*-type cytochromes revealed a decrease in *c* and *a* type cytochromes under continuous exposure to CuSO_4_ (500 - 1500 μM). In contrast, no significant change in cytochrome levels was observed following an 8 h CuSO_4_ shock in cells grown overnight in LB medium (**B**).

### Cu excess depletes *c*-type cytochromes in the cyanobacterium *Synechocystis* PCC 6803

In photosynthetic organisms, the copper-containing protein plastocyanin and the heme protein cytochrome *c*_6_ can functionally substitute for one another as electron carriers between the cytochrome *b_6_f* complex and photosystem I [42]. Under Cu limitation, green algae and cyanobacteria replace the Cu-containing plastocyanin with *c*-type cytochrome *c_6_* thereby maintaining photosynthetic electron transfer while conserving Cu for other cuproproteins. Conversely, excess Cu disrupts photosynthetic electron transport and promotes photoinhibition. Although these studies do not directly demonstrate that excess Cu inhibits cytochrome *c* assembly in thylakoids membranes, they indicate that cytochrome dependent electron transfer is sensitive to Cu imbalance. Consistent with this sensitivity, Cu sulfate has long been used to control cyanobacterial blooms and algal biomass in freshwater systems [43]. These observations prompted us to ask whether Cu excess similarly affects cytochrome *c*-dependent proteins in cyanobacteria. To address this question, *Synechocystis* PCC 6803 cells we grown in the presence of increasing CuSO_4_ concentrations (0.3-7.5 μM) and equal amounts of total protein extract were analyzed by ECL to detect *c*-type cytochromes. Excess copper progressively impaired cell pigmentation and growth (Fig. S3). At low CuSO_4_ concentrations (0.3 – 2.5 μM), the abundance of *c*-type cytochromes remained largely unchanged, whereas higher concentrations (5 and 7.5 μM) caused a marked decrease in all detected *c*-type cytochromes, including the cytochrome *b_6_f* subunits and cyt *c*_6_. Thus excess copper depletes *c*-type cytochromes also in *Synechocystis*, extending our observations in heterotrophic bacteria to a photosynthetic cyanobacterium.

### Evidence from heterologous expression in *E. coli* supports Cu-dependent disruption of cytochrome *c*-type assembly

Cytochrome *c*-type proteins can be efficiently produced in *E. coli* strains co-expressing the *ccm* gene cluster required for cytochrome c maturation [44]. In addition, excess copper has been shown to promote the formation of non-native disulfide bonds in the periplasm of *E. coli* strains lacking the periplasmic disulfide isomerase DsbC [24]. Because cytochrome *c* assembly occurs in the periplasm and requires the coordinated assembly of heme and apocytochrome components, we hypothesized that copper overload may interfere with this process. To test this hypothesis, we examined the effect of CuSO_4_ supplementation on cytochrome *c* production in *E. coli* BL21 harbouring the pEC86 plasmid, which express the *ccm* gene cluster [44]. In this strain, we expressed three proteins from *R. gelatinosus*: a soluble, truncated form of the *cbb_3_* type cytochrome *c* subunit CcoO (CcoO-H_6_), the soluble periplasmic cytochrome *c* peroxidase CcpA-H_6_, and the periplasmic dithiol: disulfide oxidoreductase DsbA-H_6_ as a negative control. These strains were grown in LB medium containing increasing CuSO_4_ concentrations and induced with IPTG. Western blot analysis of crude cell extracts from the strain expressing CcoO-H_6_ (19 kDa) revealed a signal in cells grown without CuSO_4_, whereas little or no signal was detected upon CuSO_4_ stress (**Figure 9A**).

**Figure 9.**
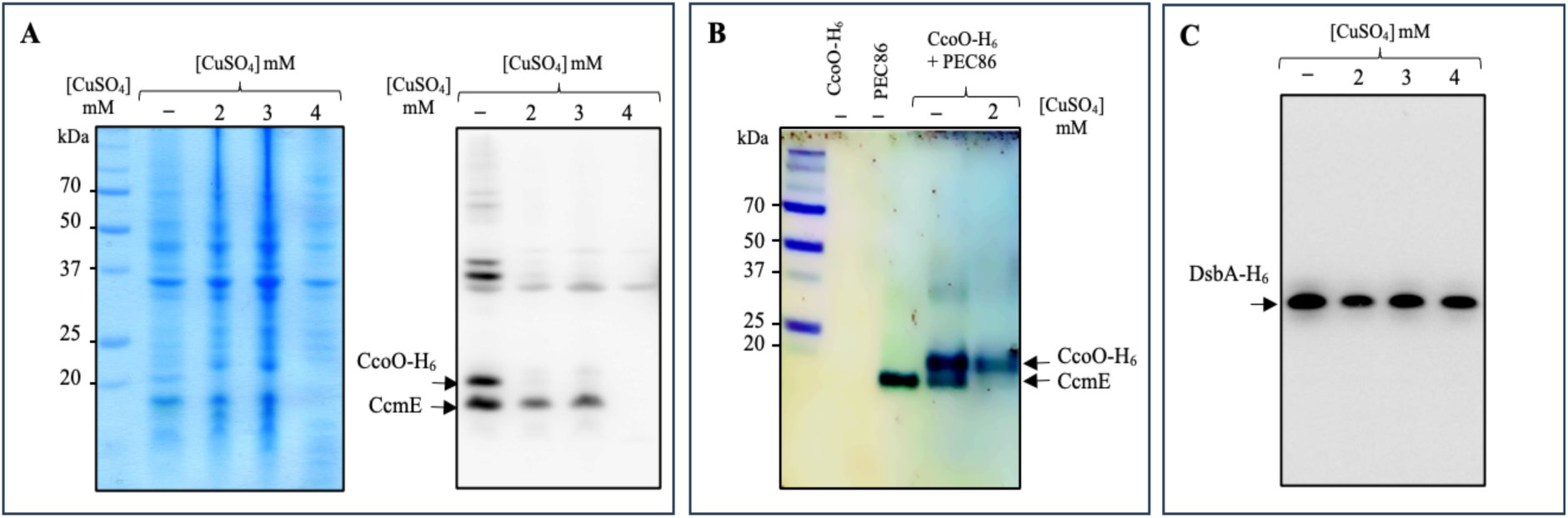
Excess Cu impairs *c*-type cytochrome synthesis in *E. coli*. Expression of the indicated proteins was analyzed in *E. coli* BL21-pEC86 cells grown in LB medium supplemented with increasing concentrations of CuSO_4_. **(A)** Coomassie-stained gel and Western blot analysis of truncated CcoO-H_6_ and CcmE. (**B**) TMBZ heme staining to assess the heme-associated signal of the expressed proteins. (**C**) Western blot showing expression of DsbA-H_6_, under the same conditions.

The abundance of CcmE (16 kDa), the heme chaperone responsible for covalent heme attachment and transfer to apocytochrome, was similarly reduced with increasing CuSO_4_ concentrations. Thus, copper excess markedly affected the accumulation of both CcmE and the apocytochrome substrate CcoO-H_6_. To directly assess heme attachment, CcoO-H_6_ was assessed by SDS-PAGE electrophoresis followed by TMBZ staining (**Fig. 9B**). In absence of CuSO_4_, two TMBZ-positive bands were observed at the expected apparent molecular masses of CcmE and CcoO-H_6_, consistent with the corresponding Western blot signals. In contrast, exposure to 2 mM CuSO_4_ markedly decreased the TMBZ signal of both proteins. Together, these data indicate that copper excess reduces the accumulation of heme associated CcmE and CcoO-H_6_ and is consistent with impaired heme incorporation during cytochrome *c*-type maturation. In contrast, the abundance of the negative control protein DsbA-H_6_, produced under identical conditions, was not affected by excess CuSO_4_, despite its multiple cysteine thiol groups that could potentially interact with Cu ions (**Fig. 9C**). A similar Cu-dependent decrease in protein abundance was also observed for CcpA-H_6_ (33 kDa) (Fig. S4). Collectively, these immunodetection and heme staining data provide strong evidence that elevated Cu levels specifically impair heme attachment and cytochrome *c*-type assembly in the periplasm.

## Discussion

In Gram-negative bacteria, the inner membrane and periplasm may be particularly vulnerable to Cu(I) toxicity [26,45]. Non-specific copper uptake can expose both compartments to excess copper, while the inner-membrane P-type ATPase CopA exports cytoplasmic Cu(I) into the periplasm [46]. Given its reactivity with protein thiols and redox processes, Cu(I) must be tightly controlled in the periplasm to maintain membrane and periplasmic integrity. Periplasmic Cu-oxidases, such as CueO from *E. coli*, together with other copper-handling systems, including CopI, PcoA and CutO, can consequently be viewed as a first line of defence and molecular sentinels that limit the accumulation of toxic Cu(I) and protect periplasmic and membrane-associated processes [16,26,30,47]. Our findings provide strong support for the particular vulnerability of this extracytoplasmic compartment to Cu(I) overload, and specifically implicate *c*-type cytochromes as a key point of fragility. The periplasm and membrane harbour numerous proteins involved in respiration, photosynthetic electron transfer, protein folding, and redox homeostasis, placing these processes in close proximity to potentially toxic Cu(I). This is especially relevant to *c*-type cytochromes, whose maturation occurs in the periplasm and depends on a precisely controlled thiol-disulfide redox environment [48]. Consistent with this, accumulation of periplasmic Cu(I) in the ΔcopI mutant results in a pronounced loss of both periplasmic and membrane-associated *c*-type cytochromes [26]. Similarly, deletion of the periplasmic multicopper oxidase CutO in *Rhodobacter capsulatus* increases Cu(I) sensitivity and compromises *cbb_3_*-type cytochrome *c* oxidase biogenesis [47]. Importantly, this vulnerability is not restricted to mutants defective in Cu-homeostasis: copper overload also decreases *c*-type cytochrome levels in wild-type bacteria, including *R. gelatinosus*, with comparable effects observed in other bacteria such as *Pseudomonas, Vibrio,* and *Bacillus*. Thus, the sensitivity of *c*-type cytochromes to Cu(I) excess appears to reflect an intrinsic vulnerability of their biogenesis and/or stability, rather than simply a secondary consequence of defective copper detoxification. A potential mechanistic link may involve the periplasmic disulfide-bonding machinery. Cu(I) has been shown to promote aberrant disulfide-bond formation in *E. coli*, thereby increasing the requirement for the disulfide isomerase DsbC [24]. This is consistent with recent evidence that excess Cu(I) catalyses protein disulfide-bond formation specifically in the bacterial periplasm, and not in the cytoplasm [25]. Conversely, perturbation of the DsbA-dependent oxidative folding pathway in *R. capsulatus* compromises *cbb_3_*-type cytochrome *c* oxidase biogenesis, highlighting the importance of periplasmic redox homeostasis for cytochrome maturation [49]. Moreover, the induction of the *dsb* genes, encoding periplasmic thiol-disulfide interchange proteins, together with the striking induction (more than 2000-fold) of PA2807, the CopI homologue, in the periplasm of *P. aeruginosa* under Cu(I) excess [33], leaves little doubt as to the critical importance of protecting this compartment against the effects of Cu(I), and of safeguarding redox-dependent systems, including cytochrome *c* biogenesis. Because *c*-type cytochrome maturation requires coordinated thiol oxidation, reduction, and subsequent heme ligation, disruption of this redox environment by excess Cu(I) could interfere with multiple steps of the maturation pathway. Cu(I) may therefore compromise cytochrome *c* biogenesis either directly, through interactions with thiol-containing substrates (apocytochromes) including Ccm maturation factors, or indirectly, by disrupting the redox conditions required for proper protein folding and heme attachment. The convergence of phenotypes observed in Cu-homeostasis mutants, such as ΔcopI and ΔcutO, with those elicited by Cu challenge in WT bacteria, argues that *c*-type cytochrome biogenesis is not merely a collateral casualty of generalised Cu(I) toxicity, but represents a particularly sensitive process within the Cu-exposed extracytoplasmic compartment. We propose that the spatial coupling of Cu trafficking, redox homeostasis, and *c*-type cytochrome maturation creates a periplasmic vulnerability in which accumulation of Cu(I) can compromise cytochrome assembly. Such impairment would have profound consequences for respiratory and photosynthetic electron transfer, explaining why preservation of low, tightly controlled [50,51] levels of periplasmic Cu(I) and/or its oxidation to less toxic Cu(II) are critical for maintaining the integrity of the bacterial membrane and periplasm system.

Our findings may therefore also have implications for the control of bacterial pathogens, particularly those that thrive under microaerobic or anaerobic conditions and rely extensively on *c*-type cytochromes for respiration and ATP synthesis. Many clinically important pathogens encounter oxygen-limited environments during colonisation and infection, where alternative respiratory pathways and *c*-type cytochrome-containing electron-transfer systems become critical for growth and persistence. Our results raise the possibility that Cu(I) overload could exploit this metabolic dependency by selectively compromising cytochrome *c* biogenesis and function. This possibility is particularly attractive in the context of multidrug resistant (MDR) pathogens, for which alternative antimicrobial strategies are urgently needed. Copper is already recognised as a potent antimicrobial agent, with demonstrated activity against a broad range of MDR organisms [52,53], including methicillin-resistant *Staphylococcus aureus*, multidrug-resistant *Burkholderia cepacia* complex [54], and carbapenemase-producing Gram-negative pathogens [55]. Nevertheless, the molecular basis of Cu-mediated killing remains multifactorial, encompassing disruption of membranes, proteins, metal homeostasis, and redox balance. Inhibition of *c*-type cytochrome biogenesis represent an additional, and potentially underappreciated, component of this toxicity, particularly under conditions in which the periplasm is exposed to elevated Cu(I). Nevertheless, the composition and flexibility of the terminal respiratory chain may represent an important determinant of bacterial tolerance to Cu(I) stress. Bacteria that rely predominantly on *bo_3_* or *bd*-type quinol oxidases, or on *aa_3_*-type cytochrome oxidase, may be better able to preserve respiratory chain function and thus exhibit greater resistance to excess Cu(I). This tolerance likely reflects differences in the intrinsic susceptibility of individual respiratory components to Cu-mediated damage, together with the capacity of these alternative oxidases to sustain electron transport when other, more Cu-sensitive components of the chain are compromised. *P. aeruginosa* illustrates this principle directly. In contrast to the *cbb_3_* complex, the *aa_3_* oxidase lacks *c*-type cytochrome, rendering it inherently less susceptible to Cu(I) mediated damage. The coexistence of both oxidases within the same organism therefore provides a built-in respiratory safety net: even when Cu excess compromises *cbb_3_* biogenesis, the *aa_3_* branch can in principle, sustain aerobic respiration, albeit with reduced efficiency, since the *bc_1_* complex and the soluble cytochrome *c* that feed electrons to *aa_3_* are themselves affected by Cu(I).

The second, complementary determinant of Cu tolerance is the presence of redundant and multiple periplasmic Cu-detoxification systems, again exemplified by *P. aeruginosa*. Bacteria equipped with several overlapping Cu handling systems are likely better buffered against periplasmic Cu overload than organisms relying on a single detoxification pathway (*V. cholerae*), providing a further layer of resistance that complements respiratory chain flexibility. Together, these observations suggest that bacterial resilience to Cu stress reflects the combined output of two largely independent safeguards: a respiratory chain capable of bypassing Cu-sensitive, cytochrome *c* dependent branches, and a periplasmic detoxification network sufficiently redundant and efficient to buffer against Cu overload before it reaches damaging levels. An important implication of this model is that Cu toxicity could be potentiated by disabling either safeguard, but particularly the latter, since it acts upstream to prevent Cu(I) accumulation in the first place. Genetic or pharmacological disruption of periplasmic Cu(I) detoxification systems, such as CopI, CutO, CueO, or Csp in *N. gonorrhoeae*, which act as protective barriers maintaining periplasmic Cu homeostasis, could therefore enhance the antimicrobial activity of Cu and potentially reduce the amount required to achieve bactericidal effects. This principle is well illustrated by Csp, the periplasmic Cu storage protein of *N. gonorrhoeae*, which has been shown to participate in periplasmic Cu homeostasis [56]. A *csp* deletion strain displays significantly increased susceptibility to Cu-mediated toxicity, indicating that Csp normally allows *Neisseria* to withstand Cu toxicity during infection. Consistent with this protective role, *N. gonorrhoeae* vaccine raised against Csp elicited anti-Csp antibodies in mice that showed efficient bactericidal activity against several *N. gonorrhoeae* strains [56,57]. By neutralising Csp function, such a vaccine could therefore act synergistically with the host’s natural Cu-based defences, sensitising the pathogen to host-derived Cu and limiting bacterial development during infection. The high dependence on *cbb_3_* type oxidase and on *c*-type cytochromes [10,58], together with the protective role of Csp in *N. gonorrhoeae*, identify two related vulnerabilities in this bacterium, biogenesis of *c*-type cytochromes and periplasmic Cu(I) detoxification, that could be jointly exploited to potentiate the antimicrobial activity of copper.

## Conclusions

Copper is an essential micronutrient but becomes toxic at elevated concentrations owing to its high chemical reactivity and strong affinity for thiol containing ligands. Our study identifies *c*-type cytochrome biogenesis as a conserved target of Cu toxicity across diverse bacterial species, linking Cu-mediated antibacterial activity to respiratory vulnerability. In particular, dependence on cytochrome *c*, especially the *cbb_3_*-type cytochrome *c* oxidase, may be a key determinant of Cu susceptibility. These findings have important clinical implications, suggesting that multidrug-resistant pathogens adapted to microaerobic conditions through *cbb_3_*-type oxidase activity may be particularly vulnerable to Cu excess in the periplasm, revealing a potential avenue for antimicrobial strategies.

## Materials and methods

### Bacterial strains and growth media

Strains are listed Table S1. *E. coli, B. subtillis, Vibrio cholorea, P. aeruginosa* and and *P. syringae* were grown microaerobically (50 ml flasks filled with 50 ml medium) at 37°C in LB medium. *B. japonicum* was grown microaerobically in yeast mannitol (YM). *R. gelatinosus* was grown in malate growth medium (MM) at 30°C, microaerobically in the dark (50 ml flasks filled with 50 ml medium) (30). *Synechocystis* PCC 6803 was grown photoautotrophically (2500 lux) in BG11 medium. Kanamycin, was used at 50 µg/ml for ΔcopI mutants and Ampicillin was used at 100 µg/ml and chloramphenicol at 10 µg/ml for *E. coli*. The malate medium copper concentration is 1.6 µM, and exogenous CuSO_4_, was added to the growth medium at the indicated concentrations. Bacterial strains are listed in Table S1 in the supplemental material.

### Membrane and soluble protein preparation

Cells were harvested and disrupted using a French press in 0.1 M sodium phosphate buffer (pH 7.4) containing 1 mM phenylmethylsulfonyl fluoride (PMSF). Membrane and soluble protein fractions were subsequently separated by differential ultracentrifugations as previously described [26]. The resulting pellet (membranes) was resuspended in the same buffer without PMSF. Protein concentrations were determined using the bicinchoninic acid assay (Sigma) using bovine serum albumin as the standard.

### Blue-native gel electrophoresis and *in-gel* activities

Blue-native polyacrylamide gel (BN-PAGE) electrophoresis of membrane proteins solubilized with β-dodecyl-maltoside (DDM) and subsequently *in-gel* Cox activity assays were performed as described [29,59]. Succinate dehydrogenase in-gel activity assays were performed as described for the succinate-NBT (Nitroblue tetrazolium) reductase assay in [59].

### Plasmids and expression in *E. coli*

Plasmids pET22-CcoO-H_6_, pET22-CcpA-H_6_ or pET22-DsbA-H_6_. were generated by cloning PCR products digested by NcoI and XhoI in the corresponding restriction sites within pET22-b (Novagen). Primers used are listed in Table S2. This strategy generated proteins containing the PelB signal peptide for periplasmic localization and a C-ter 6-histidines tag for immunodetection. For protein production, the pET22-derived vectors were cotransformed with the plasmid pEC86 in *E. coli* BL21. Cultures were grown aerobically in LB medium with increasing CuSO_4_ to mid-exponential phase and induced with 1 mM isopropyl β-D-thiogalactoside over-night at 30°C. Cells were harvested and pellets resuspended in 50 mM Tris-HCl, pH 7.4, 250 mM NaCl. Cells were disrupted by French-Press and protein concentrations were determined using the bicinchoninic acid assay (Sigma) using bovine serum albumin as standard.

### Western blotting and immunodetection

Twenty micrograms of total protein was separated by SDS-PAGE (12% or 14% polyacrylamide) and subsequently transferred onto Hybond ECL polyvinylidene difluoride (PVDF) membranes (GE Healthcare). His_6_-tag proteins were detected using HisProbe-HRP (Pierce) according to the manufacturer’s instructions. Reactive bands were detected using a chemiluminescent HRP substrate according to the method of Haan and Behrmann [60]. Image capture was performed with a LAS-3000 cooled charge-coupled device (CCD) camera system (Fuji).

### *In-gel* heme staining

Detection of cytochrome *c* on non-reducing SDS-PAGE was performed using 3,3′,5,5′-tetramethylbenzidine (TMBZ) as described by Thomas et al. [61]. For cytochromes of *Synechocystis* PCC 6803 cytochromes, TMBZ staining was unsuccessful (likely because of their low abundance) proteins were transferred onto PVDF membrane, and then incubated with ECL (SuperSignal^TM^ West Pico, Thermo Scientific). Image capture was performed with the LAS-3000 system.

### Spectrophotometric measurements

Absorption spectroscopy was performed with a Cary 500 spectrophotometer. For reduced minus oxidized difference spectra, total pigments were extracted from cell pellets or membrane fractions with acetone-methanol (7/2 [vol/vol]). For each sample, an initial spectrum was collected on oxidized sample upon addition of 50 μM K_6_Fe(CN)_3_. The sample was then reduced by adding a few crystals of dithionite, and a second spectrum collected. The reduced-minus-oxidized difference spectrum was subsequently calculated from the two spectra.

## Acknowledgments

The authors thank Dr. Christine Pourcel, Dr. Yamaichi Yoshiharu, Dr. Peter Maerget, from I2BC and Dr. Cécile Raynaud from IPS2 for providing *P. aeruginosa, V. cholerae, B. japonicum,* and *P. syringae*, respectively.

## Author contributions

- Conceptualization: Durand A., Ouchane S.
- Investigation: Durand A., Steunou A-S, Khan M-A., Bourbon M-L, and Ouchane S.
- Formal analysis: Durand A., Steunou A-S, Khan M-A, Bourbon M-L, Cassier-Chauvat C., Chauvat F., and Ouchane S.
- Funding acquisition: Ouchane S.
- Project administration: Durand A. Ouchane S.
- Supervision: Ouchane S.
- Writing original draft: Ouchane S.
- Writing review & editing: Durand A., Steunou A-S, Khan M-A, Cassier-Chauvat C., Chauvat F., and Ouchane S.

## Data availability statement

Data supporting the findings of this work are available within the paper and its Supplementary Files. The datasets materials generated and analyzed in the study are available from the corresponding authors upon request.

## Declaration of generative AI assisted technologies

The authors used AI for language editing purposes only, then reviewed and edited the content as needed and take full responsibility for the content of the publication.

