## Supplementary material for "Copper-Mediated Antimicrobial Activity : Targeting Cytochrome *c* Complexes Across Bacterial Species": Suppl file

### Supplementary information

**Running Title**: *Respiratory Vulnerability to Copper in Bacteria*

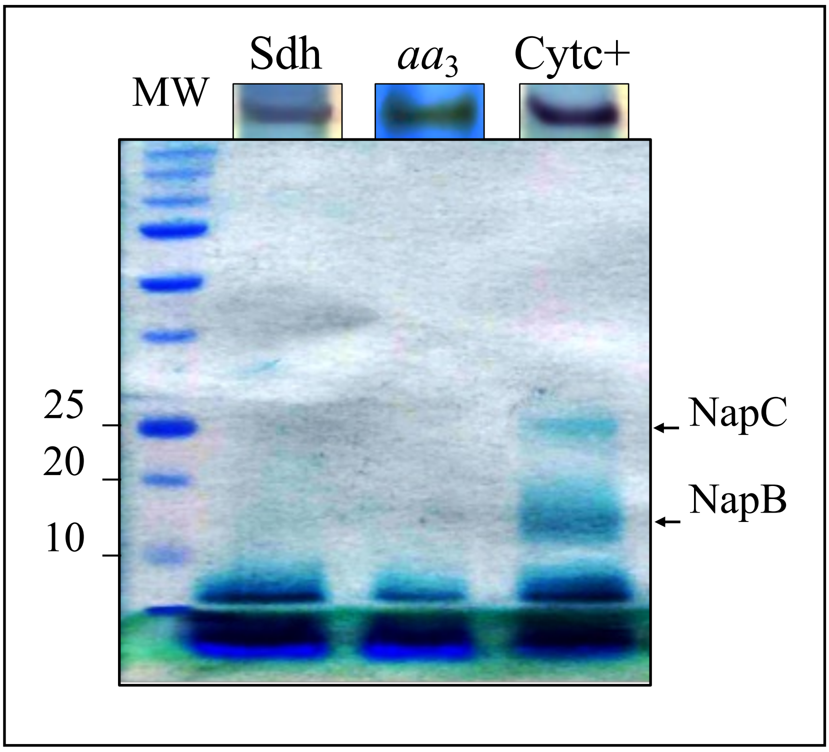

**Figure S1 : NapB (1**5 kDa **) and NapC (**25 kDa **) detection in the Cytc**⁺ **active band**: Cytc⁺ active band was excised from the stained BN-PAGE, resolved in a second dimension by SDS-PAGE followed by TMBZ staining for cyt*c* detection. The SDH and $aa_3$ bands were analyzed in parallel as negative staining controls.

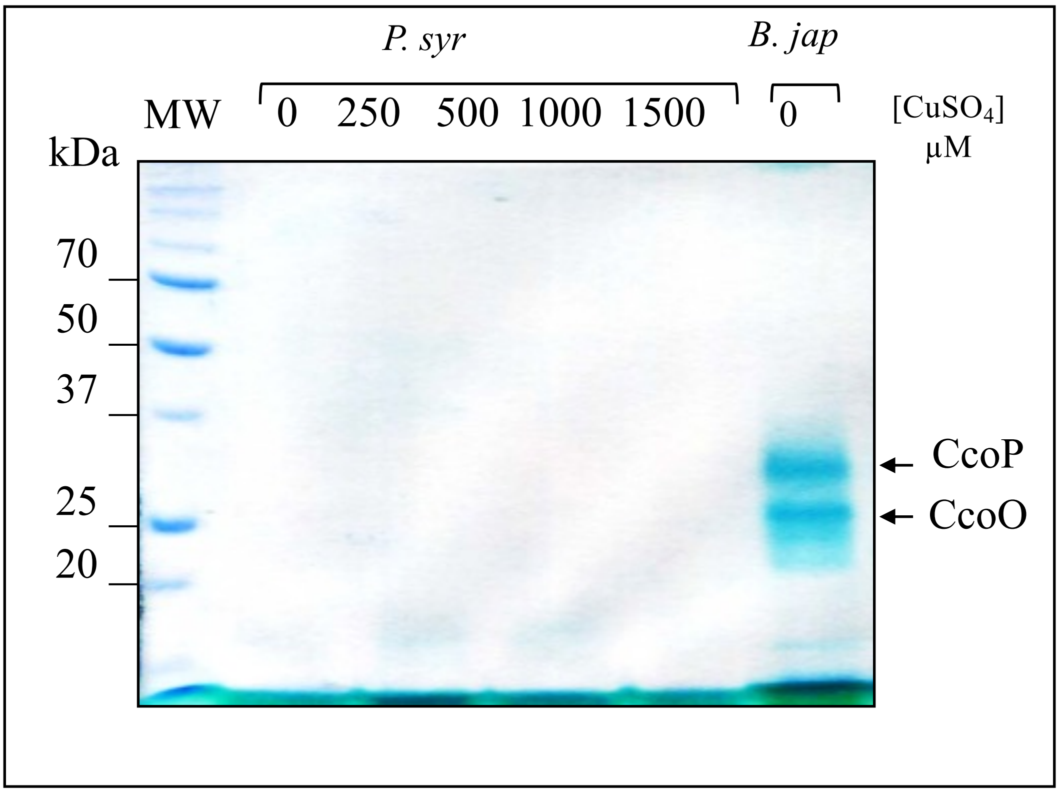

**Figure S2 :** ***Pseudomonas syringae*** **cytochrome** ***b*** **complexes withstand Cu stress that disrupts** ***c*****-type cytochromes.** *c*-type cytochrome profiles in *P. Syringae* and *B. japonicum* (positive control) by heme staining. No *c*-type was detected in P. syringae because of a non-functional CCM machinery in this strain.

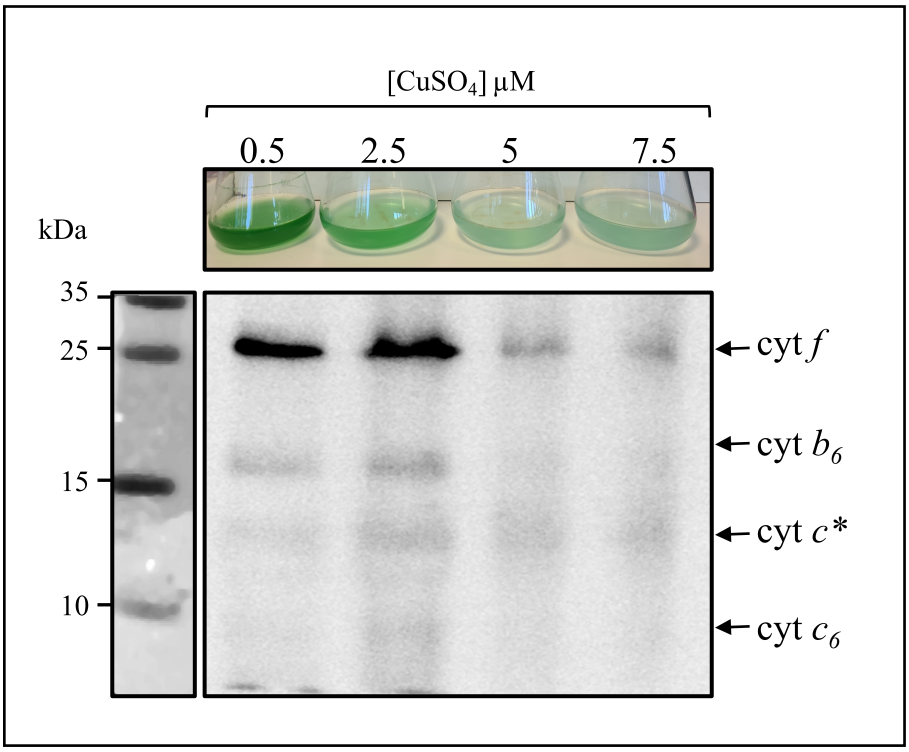

**Figure S3. Effect of $CuSO_4$ on growth and *c*-type cytochromes in *Synechocystis* PCC 6803.** *Synechosystis* PCC 6803 was grown in BG11 medium supplemented with increasing concentrations $CuSO_4$ of (2.5 - 7.5 µM). Differences in growth and pigmentation were assessed under the different copper conditions. *c*-type cytochrome profiles in total proteins fractions were by 12% SDS-PAGE and visualization with enhanced chemiluminescence (ECL). Copper (5 to 7.5 µM) supplementation altered the signals of all detected cytochromes.

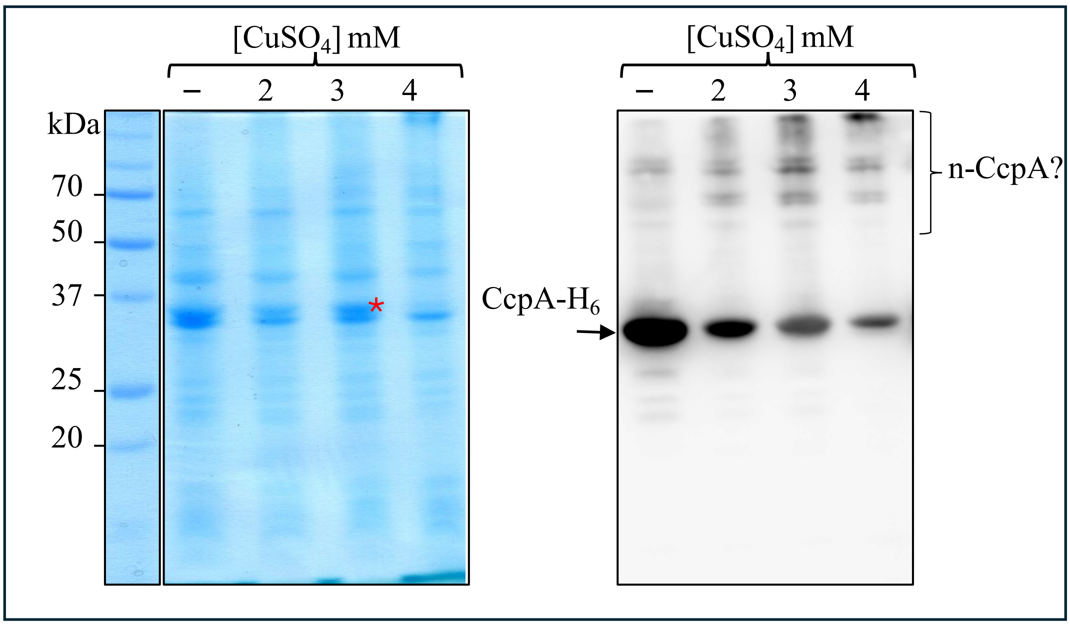

**Figure S4. Excess Cu impairs *c*-type cytochrome synthesis in *E. coli*.** Expression of CcpA-$H_6$ in *E. coli* BL21-pEC86 cells grown in LB medium supplemented with increasing concentrations of $CuSO_4$. Coomassie-stained gel and Western blot analysis showing the decrease of CcpA-$H_6$. n-CcpA corespond to putative Cu induced aggregates of CcpA-$H_6$.

**Table S1: strains and plasmids used in this study**

| | Strain designation | References |
|---|---|---|
| *E. coli* | K12 | New England Biolabs |
| *Vibrio Cholerae* | O1 El Tor, | [1] |
| *Vibrio Cholerae* | copI::Tn | [1] |
| *Pseudomonas aeruginosa* | PAO1 | [2] |
| *Pseudomonas syringae* | pv. tomato DC3000 | [3] |
| *Bradyrizobium japonicum* | USDA 110 | [4] |
| *Bacillus subtilis* | 168 | [5] |
| *E. coli* | BL21 (DE3) | New England Biolabs |
| *Rubrivivax gelatinosus* | Strain S1 | [6] |
| *Synechocystis sp.* | PCC 6803 | Pasteur institute (Paris, France) |
| **Plasmids** | | |
| pET22b | Protein expression plasmid | Novagen |
| pEC68 | *ccm* gene cluster containig plasmid, | [7] |
| pET22-CcoO-$H_6$ | *ccoO* PCR product cloned in pET22b at NcoI and XhoI sites | This study |
| pET22-CcpA-$H_6$ | *ccpA* PCR product cloned in pET22b at NcoI and XhoI sites | This study |
| pET22-DsbA-$H_6$ | *dsbA* PCR product cloned in pET22b at NcoI and XhoI sites | This study |

**Table S2 : primers used in this study**

| Primer | 5'-3' Sequence |
| --- | --- |
| ccoO-NcoI _Fw | TGGCCATGGTGTTCTTCCAGCGCTCCAC |
| ccoO-XhoI _Rv | GGTGCTCGAGCTTGACGGCCGTGCCCAGC |
| ccpA-NcoI _Fw | GGCCATGGCCCAGAGCCAGGAGCCGATCC |
| ccpA-XhoI _Rv | TGGTGCTCGAGCTTGAACGGATGCGGACG |
| dsbA-NcoI _Fw | ATGGCCATGGCGCAGGGCGGGCCGGTCGA |
| dsbA-XhoI _Rv | GTGCTCGAGGGCGCCCTGGCGGGCGCGGCC |
